# Brain-wide mapping of neuroanatomical connections to the auditory cortex of hearing and deaf mice

**DOI:** 10.64898/2026.04.15.718225

**Authors:** Thomas C. Harmon, Andrew Jin, Evelyn J. Hardin, Richard Mooney

## Abstract

Remarkable therapeutic innovations have made it possible to establish hearing in congenitally deaf subjects. Despite these advances, a potential obstacle to restoring auditory function is that the absence of auditory experience alters the connectivity of the auditory cortex, a region that contributes to auditory perception and cognition. Here we used an intersectional genetic approach to map the brainwide inputs to the primary auditory cortex of congenitally deaf mice and their hearing littermates. We found that deaf mice displayed a significant reduction in afferents arising from the basomedial amygdala, the core of the medial geniculate nucleus, and anterior auditory thalamic nuclei. Nonetheless, major aspects of auditory cortical connectivity, including input from other thalamic nuclei and from non-auditory regions of the cortex, were unaffected by deafness. These findings highlight altered and preserved connectivity of the auditory cortex in the absence of auditory experience, which may inform therapies designed to establish hearing in congenitally deaf subjects.

## Introduction

The auditory cortex is crucial to the perception of complex sounds, including speech, and its functional properties are highly sensitive to auditory experience, including the effects of hearing loss. Intriguingly, many forms of congenital deafness arise from genetic mutations that impair cochlear hair cell function, but that leave the afferent pathway to the central auditory system intact (Akil et al., 2012; Kawashima et al., 2011; Nist-Lund et al., 2019; Roux et al., 2006). An important question addressed here is how genetic insults that selectively impact hair cells and produce congenital deafness alter higher regions of the central auditory system, particularly the auditory cortex.

The connectivity of the auditory cortex has been extensively studied using a wide variety of anatomical tracing methods in a variety of adult mammals with normal hearing, including mice (da Costa et al., 2017; Pan et al., 2025). Indeed, the primary auditory cortex is a site of remarkable convergence of input from the auditory thalamus, other cortical regions – including somatosensory, visual, premotor, associational, and secondary auditory areas – and the cortical subplate, including the amygdala. This remarkable degree of convergence enables the auditory cortex to integrate bottom-up signals conveying auditory information with top-down signals important to other forms of sensory processing, movement, motor planning, and decision making (e.g. Atilgan et al., 2018; Francis et al., 2018; Gale et al., 2021; Iurilli et al., 2012; Lohse et al., 2021; Napoli et al., 2021; Nelson et al., 2013; Schneider et al., 2014). Indeed, the integration of these auditory and non-auditory signals is clearly evident during vocalization, where motor-related signals suppress vocalization-related auditory feedback (Eliades & Wang, 2003; Müller-Preuss & Ploog, 1981; Numminen & Curio, 1999). Moreover, in mice with congenital sensorineural deafness, non-auditory signals generated during vocalization not only persist but can be enhanced (Harmon et al., 2024). Furthermore, visual and somatosensory responses are enhanced in the auditory cortex of deaf mice (Hunt et al., 2006), as well as in the auditory cortices of deaf ferrets, cats, and humans (Allman et al., 2009; Auer et al., 2007; Finney et al., 2001; Karns et al., 2012; Meredith & Allman, 2012). However, whether such enhancement reflects structural reorganization of auditory cortical afferents, or functional modification of an otherwise stable, experience-independent afferent architecture, is not fully resolved.

In fact, numerous studies in a variety of mammalian species have explored how early hearing loss alters auditory cortical connectivity. For example, studies using conventional retrograde tracers in white cats, which display profound hearing loss early in postnatal life, suggest that the gross connectivity of the auditory cortex can be established and maintained without auditory experience. Nonetheless, these studies also revealed that, in deaf subjects, the primary auditory cortex receives reduced auditory thalamic input and ectopic visual thalamic input, while secondary auditory cortex received expanded visual and somatosensory cortical input (Barone et al., 2013; Chabot et al., 2015; Kok et al., 2014; Kral et al., 2003; Wong et al., 2014). In contrast, the pattern and number of inputs to the primary auditory cortex is highly similar in ferrets subjected to ototoxic lesions early in postnatal life when compared to those with normal hearing (Meredith & Allman, 2012). However, the genetic mutation that gives rise to deafness in white cats exerts broader effects on the auditory periphery compared to pointillistic mutations, such as Tmc1^Δ/Δ^ (Kawashima et al., 2011), that specifically block mechanoelectrical transduction by cochlear hair cells, and ototoxic lesions made early in postnatal life may not accurately model congenital deafness in human subjects. Moreover, while conventional tracers such as dextrans are useful, intersectional methods that use retrograde viruses in transgenic reporter mice afford a degree of sensitivity and brain-wide neural circuit mapping that is especially well-suited to analyze the organization of auditory cortical afferents.

To better understand how congenital sensorineural deafness impacts the structural organization of auditory cortical afferents, we injected retrograde viral vectors for Cre-recombinase into the auditory cortex of adult deaf (Tmc1^Δ/Δ^) and hearing (Tmc1^+/Δ^) mice that had been crossed into a Cre-recombinase reporter line (Ai14). This approach resulted in widespread labeling of auditory cortical afferents across cortical and subcortical regions of both deaf and hearing mice. Deaf mice displayed a significant reduction in afferents arising from the basomedial amygdala, the core of the medial geniculate nucleus, and anterior auditory thalamic nuclei. Nonetheless, major aspects of auditory cortical connectivity, including input from other thalamic nuclei and from non-auditory regions of the cortex, were unaffected by deafness. Therefore, congenital deafness results in a landscape of structural alteration and preservation that interventional therapies for hearing restoration can exploit.

## Results

### Brain-wide mapping of auditory cortical afferents in hearing and deaf mice

To identify auditory cortical afferent neurons, we first crossed Cre-dependent tdTomato reporter mice (Ai14) with transgenic mice with a nullifying mutation to the TMC1 locus (Tmc1^Δ/Δ^; Kawashima et al., 2011). We backcrossed double heterozygous offspring to produce litters of hearing Tmc1^+/Δ^ and deaf Tmc1^Δ/Δ^ mice (Figure 1A). In adult Tmc1^+/Δ^ but not Tmc1^Δ/Δ^ mice, recordings of auditory brainstem responses to brief clicks elicited a multicomponent response indicative of hearing (Figure 1B). We co-housed mice with same-sex littermates and matched the experimental schedule for all mice within a litter. This design ensured that hearing and deaf littermates had comparable non-auditory experiences and neurodevelopmental maturity at the time that auditory cortical afferent neurons were labeled.

**Figure 1:**
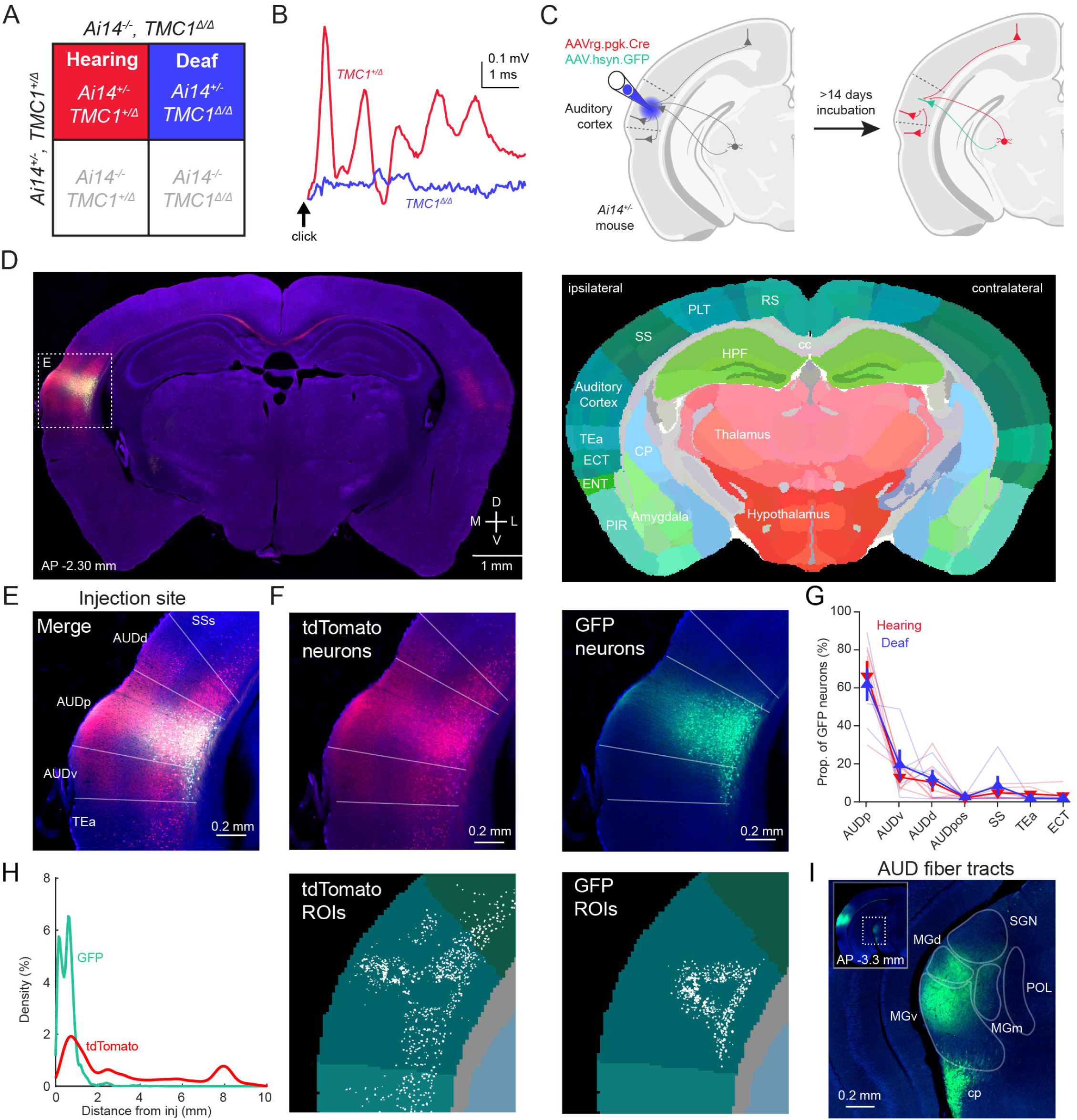
Brain-wide labeling of auditory cortical afferent neurons. **A)** Punnett square of genotypes resulting from crossing hemizygous Ai14+/-/Tmc1+/A mouse with Ai14-/-/Tmc1A/A mouse. Colored squares, genotypes of mice used in the experiment. **B)** Sample recordings, collected from either a Tmc1+/A (red) or Tmc1 A/A (blue) mouse, of auditory brainstem responses to brief clicks. **C)** Schematic of viral strategy for labeling auditory cortical afferent neurons and identifying the injection site. **D)** Left, example section collected near the injection site from the brain of a deaf mouse. Right, custom atlas projection generated for this section by the NUTIL software. **E)** Magnified image of the injection site in D with all color channels merged. **F)** Top, image in E with color channels separated to show tdTomato-expressing neurons (left) and GFP-expressing neurons (right), contrasted with blue fluorescent nuclear staining. Bottom, projections of the brain regions (colored areas) and identified ROIs (white masks) corresponding to top images. **G)** Mean ± SEM of the proportion of GFP neurons assigned to regions surrounding the injection site, calculated across all hearing (red) or deaf (deaf) mice. **H)** Kernel-smoothed density plot of the euclidean distance of GFP (green) or tdTomato (red) ROIs from the injection site. **I)** Magnified image of GFP-labeled axons in the medial geniculate nuclei or the cerebral peduncle. Inset, ipsilateral hemisphere of section showing source of the magnified image (dashed line box). Abbr: AUDd, AUDp, AUDv - dorsal, primary, and ventral auditory cortex; cc - corpus callosum; CP - caudoputamen; cp - cerebral peduncle; EOT - ectorhinal cortex; ENT - entorhinal cortex; HPF - hippocampal formation; MGd, MGm, and MGv - dorsal, medial, and ventral part of the medial geniculate nucleus; PIR - piriform cortex; POL - posterior limiting nucleus; PTL - posterior parietal cortex; RS - retrosplenial cortex; SGN - suprageniculate nucleus; SS - somatosensory cortex; TEa - temporal association.

To localize our injection site while also labeling auditory cortical afferent neurons, we injected a cocktail of AAV2/9-hsyn-eGFP and AAVrg-pgk-Cre (GFP and retro-Cre) into the left auditory cortex of the double transgenic mice above (Figure 1C). This technique resulted in GFP expression at the injection site and tdTomato labeling of afferent neurons throughout the brain, including with the GFP-delimited injection site (Figure 1D-F). We used the QUINT image analysis pipeline to assign fluorescent neurons to regions of the brain (Yates et al., 2019).

Within the pipeline, we generated custom anatomical atlas projections by aligning the Allen Institute Mouse Brain Atlas to the position and slicing angle of each brain section (Figure 1D, right). In parallel, we trained and applied an object classifier to identify regions of interest (ROIs) comprising cell bodies of fluorescent neurons (Figure 1F, bottom). We then assigned ROIs to brain regions by projecting them into the custom atlas.

While tdTomato-expressing neurons were distributed throughout the brain, nearly all GFP-labeled neurons were within 1 mm of the injection site (Figure 1H). Alignment to custom anatomical atlas projections revealed that most GFP-labeled neurons were located in the primary auditory cortex (Figure 1G), and allowed us to exclude animals in which virus spread outside of the intended target region. Importantly, the distribution of GFP-labeled neurons was similar in hearing and deaf mice (F(1,5) = 1.75, p = 0.13), indicating that the placement of our injections was the same across groups. Finally, in both hearing and deaf mice, we observed GFP-labeled axons of the corticothalamic tract in the medial geniculate nucleus and axons of the corticofugal tract in the cerebral peduncle (Figure 1I), two major projection pathways from the auditory cortex. Along with the atlas alignment, this pattern of anterograde GFP labeling confirmed that our viral injections were accurately targeted to the primary auditory cortex.

In summary, this approach combined robust fluorescent labelling of neurons with an automated and standardized analysis pipeline for assigning neurons to brain regions, allowing for regional populations of auditory cortical afferent neurons to be compared between groups of hearing and deaf mice.

### Isolating the influence of auditory experience on auditory cortical afferents

To determine whether auditory experience influences the number of neurons that provide input to the auditory cortex, we first controlled for additional factors that could affect neuron counts. One prominent factor is the efficacy of viral transfection, which can vary substantially between animals injected with the same volumes and titers of virus. Indeed, for both hearing and deaf mice, counts of afferent neurons labeled through transfection of the retro-Cre vector ranged by an order of magnitude across brains (Figure 2A). We reasoned that, because suspensions of retro-Cre and GFP vectors were mixed and co-injected, counts of GFP-labeled neurons in the injection site could provide an estimate of the transfection efficacy of retro-Cre virus that is independent of hearing condition. Consistent with this possibility, counts of GFP neurons covaried with counts of tdTomato neurons when collected across the brain or within individual regions (Figure 2A, 2B).

**Figure 2:**
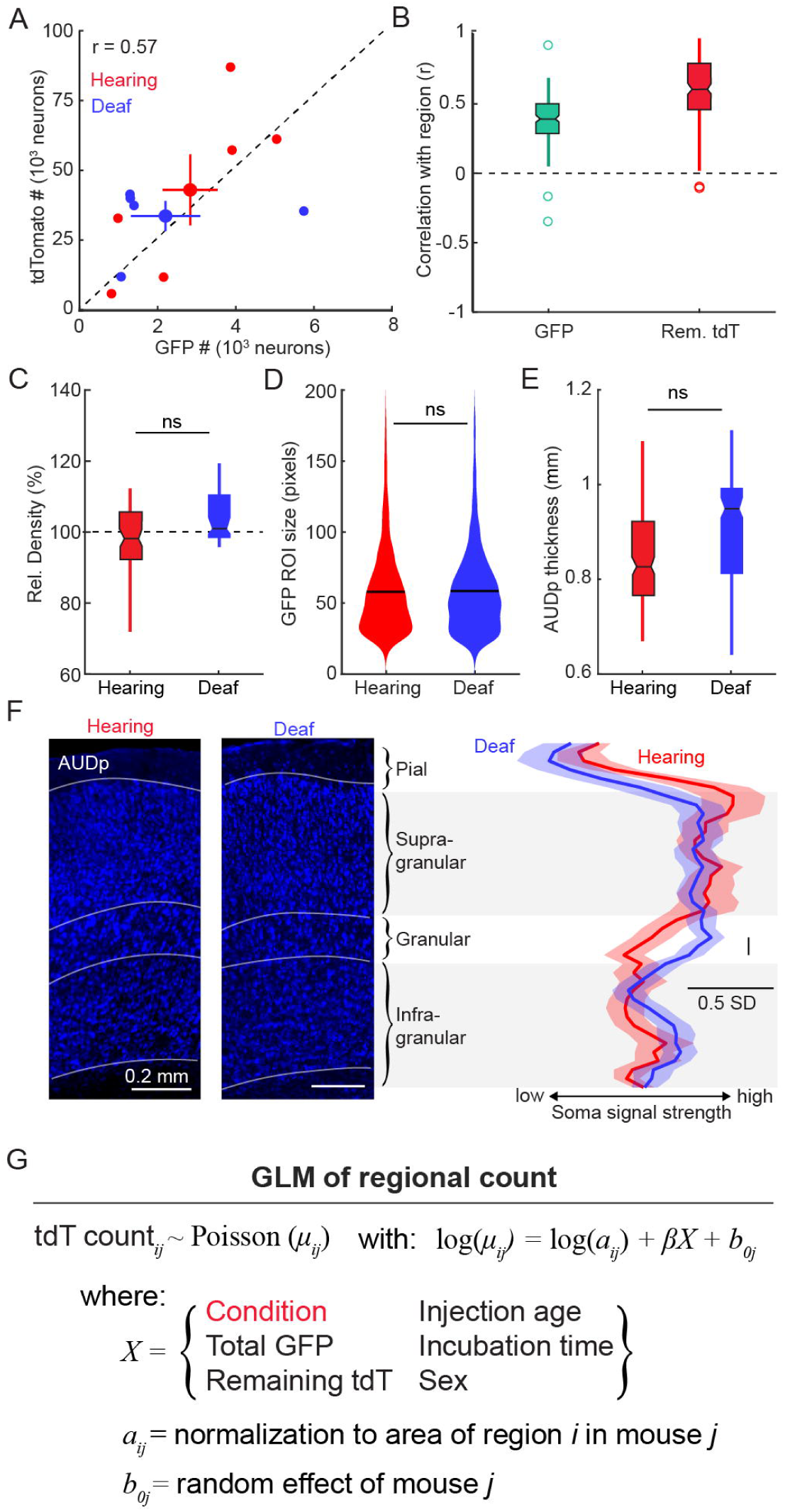
Isolating the influence of auditory experience on auditory cortical afferents. **A)** The number of GFP-labeled neurons and the number of tdTomato-labeled neurons for each brain in the dataset (small markers) and the mean ± SEM (large markers) of counts from brains of hearing (red, n = 6) or deaf (blue, n = 5) mice. Dashed line, linear fit to all data points. Pearson correlation calculated across all brains. **B)** Box and whisker plot of pearson correlation values calculated between regional tdTomato-labeled neuron counts and GFP-labeled neuron counts (green, left) or between regional tdTomato-labeled neurons and counts of tdTomato labeling in all remaining brain areas (red, right). Open circles, outliers (> 3 STD from mean). **C)** Box and whisker plot of relative cell density of the primary auditory cortex among brains from hearing (red, n = 11) or deaf (blue, n = 11) mice. **D)** Kernel smoothed density plot (bandwidth = 5) of GFP ROI size from primary auditory cortex of hearing (red, n = 5719 ROIs) or deaf (blue, n = 4398) mice. Black line, mean. **E)** Box and whisker plot of the thickness of the primary auditory cortex among brains from hearing (red, n = 11) or deaf (blue, n = 11) mice. **F)** Left, example images blue fluorescent Nissl stain from a hearing (left) and deaf (right) mouse showing the lamina of the primary auditory cortex (AUDp). Boundaries determined by visual examination of neuron size and density. Right, mean ± SEM of z-scored signal strength across the depth of the auditory cortex. Bin, 2.5% of cortical depth. *, p < 0.05. **G)** General formula of generalized linear models built for each brain region.

Counts of GFP-labeled neurons can be used to estimate transfection efficacy only if the total number of neurons is the same in the auditory cortex of deaf mice as it is in hearing mice. In the congenitally deaf cat, the thickness of the granular and infragranular layers of the auditory cortex is reduced (Berger et al., 2017), indicating that the number of neurons or the volume of their somata and dendrites is decreased. To examine whether the total number of neurons in the mouse auditory cortex was affected by deafness, we measured the optical signal strength of the blue fluorescent Nissl stain in the auditory cortex contralateral to the injection, and normalized the values to the signal strength measured elsewhere in the same section. This analysis detected no difference in the density of neurons between hearing and deaf mice (p = 0.80; Figure 2C). Furthermore, we found that the soma size of GFP-labeled neurons was indistinguishable between the groups (p = 0.34; Figure 2D).

We further compared the cytoarchitecture of the auditory cortex in hearing and deaf mice. We found no difference in the thickness of the primary auditory cortex between hearing and deaf mice (p = 0.60; Figure 2E). We also examined the density of neurons across cortical layers by measuring the normalized optical signal strength along the depth of the auditory cortex. We found that the signal strength, and therefore the density of neurons, was comparable across the infragranular, supragranular, and pial layers. Notably, we observed that the density of neurons in the granular layer was greater in deaf mice than in hearing mice (Figure 2F). In summary, the laminar structure of the primary auditory cortex is largely similar in hearing and deaf mice, indicating that counts of GFP neurons are useful to estimate transfection efficacy of retro-Cre virus across individual injections and between groups of hearing and deaf mice.

We use generalized multivariate linear models to predict regional counts of auditory cortical afferent neurons from individual brain regions to detect whether the absence of auditory experience systematically increases or decreases the number of auditory cortical afferent neurons (Figure 2G). Each model attempted to reconstruct Poisson distributions of regional neuron counts gathered from hearing and deaf mice. Within each model, we included the mouse’s hearing condition and other predictor variables that accounted for technical and biological factors that plausibly influence afferent neuron labeling. We applied this approach to counts from regions across the ipsilateral and contralateral cerebral cortex, ipsilateral cortical subplate, and ipsilateral thalamus, resulting in a systematic survey of all sources of auditory cortical afferents in hearing and deaf mice. These models returned the weighting coefficient of the hearing condition, its 95% confidence interval, and a raw p-value. While many regions displayed a non-zero coefficient, consistent with an effect of hearing condition, none of the p-values of these coefficients survived correction for multiple comparisons. Therefore, we used a qualitative criterion where an effect of hearing condition on afferent neuron labeling within a region required that the confidence interval was non-overlapping with a weighting coefficient equal to zero.

### Cortical afferents to the auditory cortex are largely unaffected by auditory experience

Our analysis revealed that the vast majority (96%) of tdTomato-expressing neurons in hearing and deaf mice were located in cortical regions, with most (86%) in the ipsilateral hemisphere. In all cortical regions we examined, afferent neurons were present in both hearing and deaf mice, indicating that, in deaf mice, no regional projections to the auditory cortex were eliminated, nor were ectopic projections established *de novo*. Ordering counts of tdTomato-labeled neurons showed that the relative numbers of auditory cortical afferent neurons in different cortical regions are similar in hearing and deaf mice (Figures 3-8). Thus, the gross organization of cortical inputs to the auditory cortex is preserved in deaf mice.

**Figure 3:**
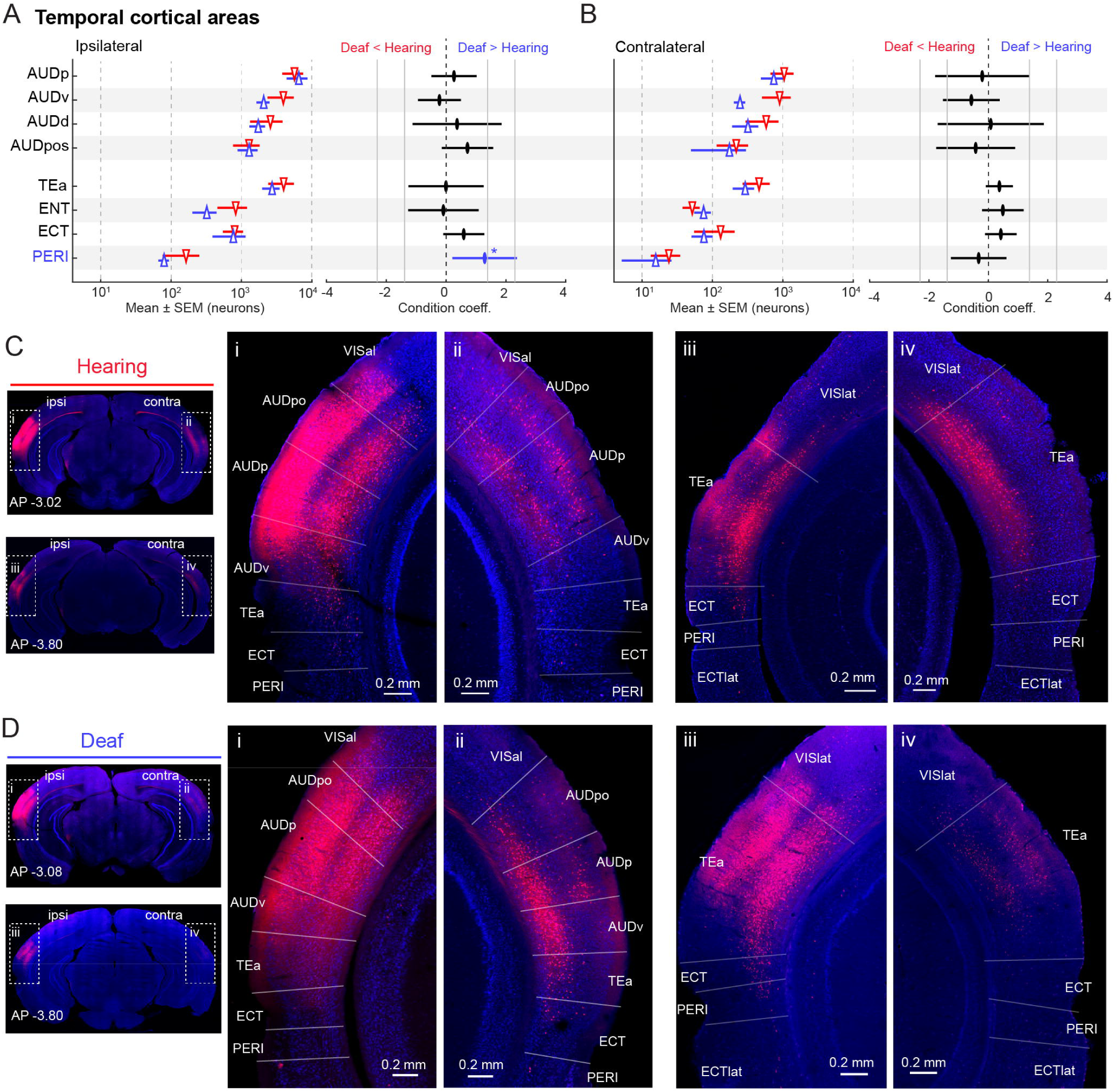
Auditory cortical afferent neuron labeling in temporal cortical areas. **A)** Left, mean ± SEM of afferent neuron counts from ipsilateral auditory regions (top) and adjacent associational cortical regions (bottom) from hearing (red) or deaf (blue) mice. Values plotted on log scale. Right, coefficient and 95% confidence interval of hearing condition for generalized linear model of regional neuron counts. Grey vertical lines, coefficient values corresponding to 4- and 10-fold difference in counts between hearing and deaf mice. *, p < 0.05. **B)** Same as A, but for contralateral regions. **C)** Examples of auditory cortical afferent neuron labeling in temporal cortical regions immediately proximal or posterior to the injection site in a hearing mouse. **D)** Same as B, but for a deaf mouse. Abbreviations: AUDp, AUDv, AUDd, and AUDpos - primary, ventral, dorsal, and posterior auditory cortical area; ECT - ectorhinal cortex; ECTIat - lateral ectorhinal cortex; ENT - entorhinal cortex; PERI - perirhinal cortex; TEa - temporal association cortex; VISIat - lateral visual cortex.

For almost all regions of the ipsi- or contralateral cerebral cortex, we did not detect an effect of auditory experience on the number of auditory cortical afferent neurons. In both hearing and deaf mice, we observed dense labeling in secondary auditory regions surrounding the primary auditory cortex, as well as in regions immediately dorsal (e.g. anterolateral visual cortex) or ventroposterior (e.g. temporal association area; Figure 3) to the injection site.

Although our model indicated a significant positive effect of deafness on labeling in the ipsilateral perirhinal cortex (p = 0.029; Figure 3A,Ciii,Diii), this effect was likely attributable to low variance in overall very sparse labeling of perirhinal cortical neurons in both hearing and deaf mice.

The numbers of tdTomato-labeled neurons in non-auditory sensory cortical regions anterior or posterior to the primary auditory cortex were also indistinguishable in hearing and deaf mice (Figure 4). Among anteriorly located regions, somatosensory areas and integrative sensory areas (e.g., agranular insula, gustatory area, and visceral area) featured sparse labeling of auditory cortical afferent neurons in both hearing and deaf mice (Figure 4C, 4D; example labeling in Figure 5Cii, 5Ciii, 5Dii, 5Diii). Posterior to the injection site, we observed labeling in visual cortical regions that was similar in hearing and deaf mice (Figure 4E-H). In somatosensory and visual cortical areas, afferent neurons were primarily located in supragranular and infragranular layers.

**Figure 4:**
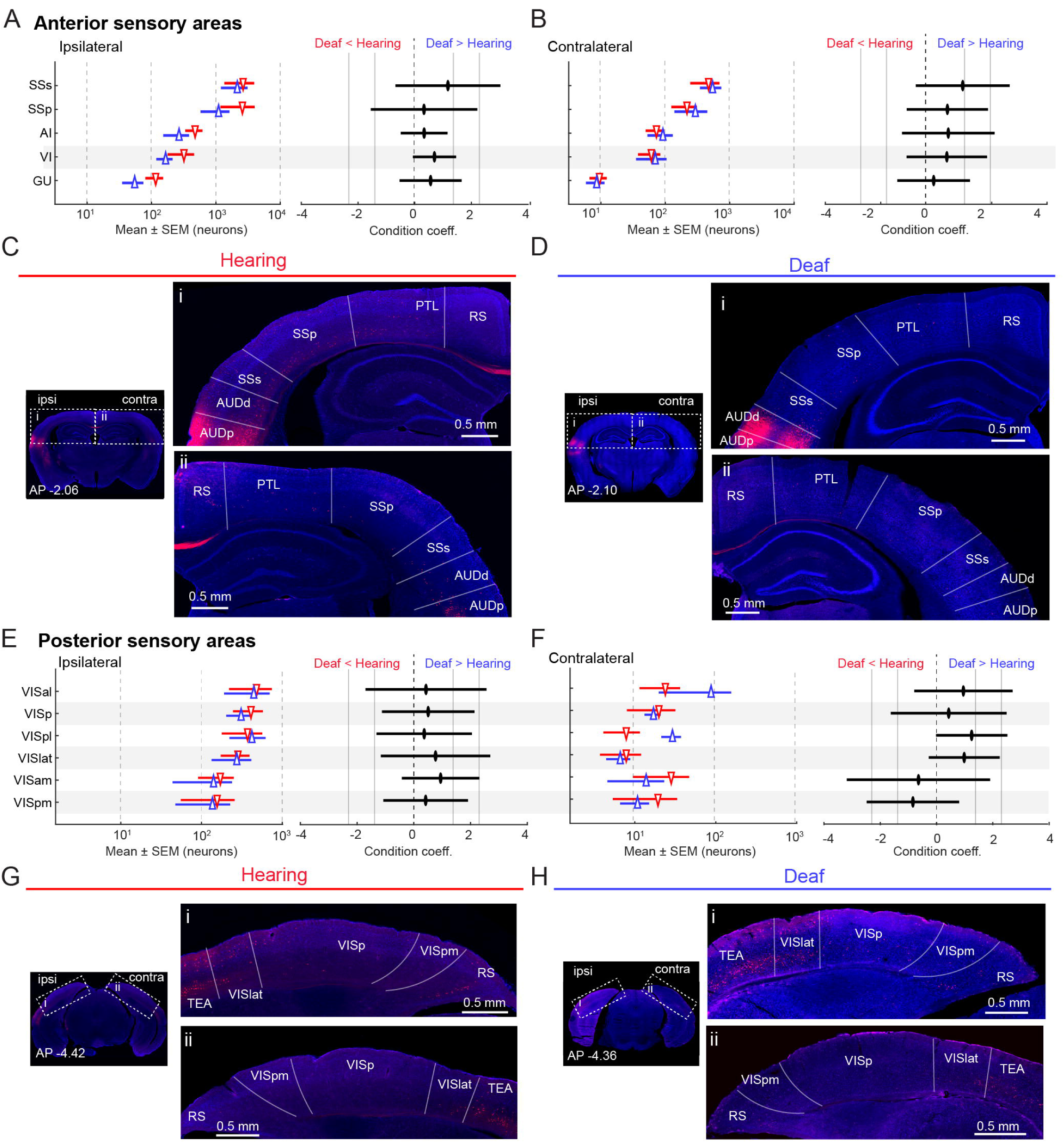
Auditory cortical afferent neuron labeling in sensory cortical areas. *A)* Left, mean ± SEM of afferent neuron counts gathered in ipsilateral sensory cortical regions located anterior to the auditory cortex in hearing (red) and deaf (blue) mice. Values plotted on log scale. Right, coefficient and 95% confidence interval of hearing condition for generalized linear model of regional neuron counts. Grey vertical lines, coefficient values corresponding to 4- and 10-fold difference in counts between hearing and deaf mice. **B)** Same as A, but for contralateral regions. **C)** Examples of auditory cortical afferent neuron labeling in sensory cortical regions located anterior to the injection site in a hearing mouse. **D)** Same as B, but for a deaf mouse. **E-H)** Same as A-D but for posteriorly-located sensory cortical regions. Abbreviations: Al - agranular insular cortex; AUDp - primary auditory cortex; AUDd - dorsal auditory cortex; GU - Gustatory cortex; SSp - primary somatosensory cortex; SSs - secondary somatosensory cortex; VI - visceral cortex; VISal, VISp, VISpI, VISIat, VISam, and VISpm - anterolateral, primary, posterolateral, lateral, anteromedial, and posteromedial visual cortex.

**Figure 5:**
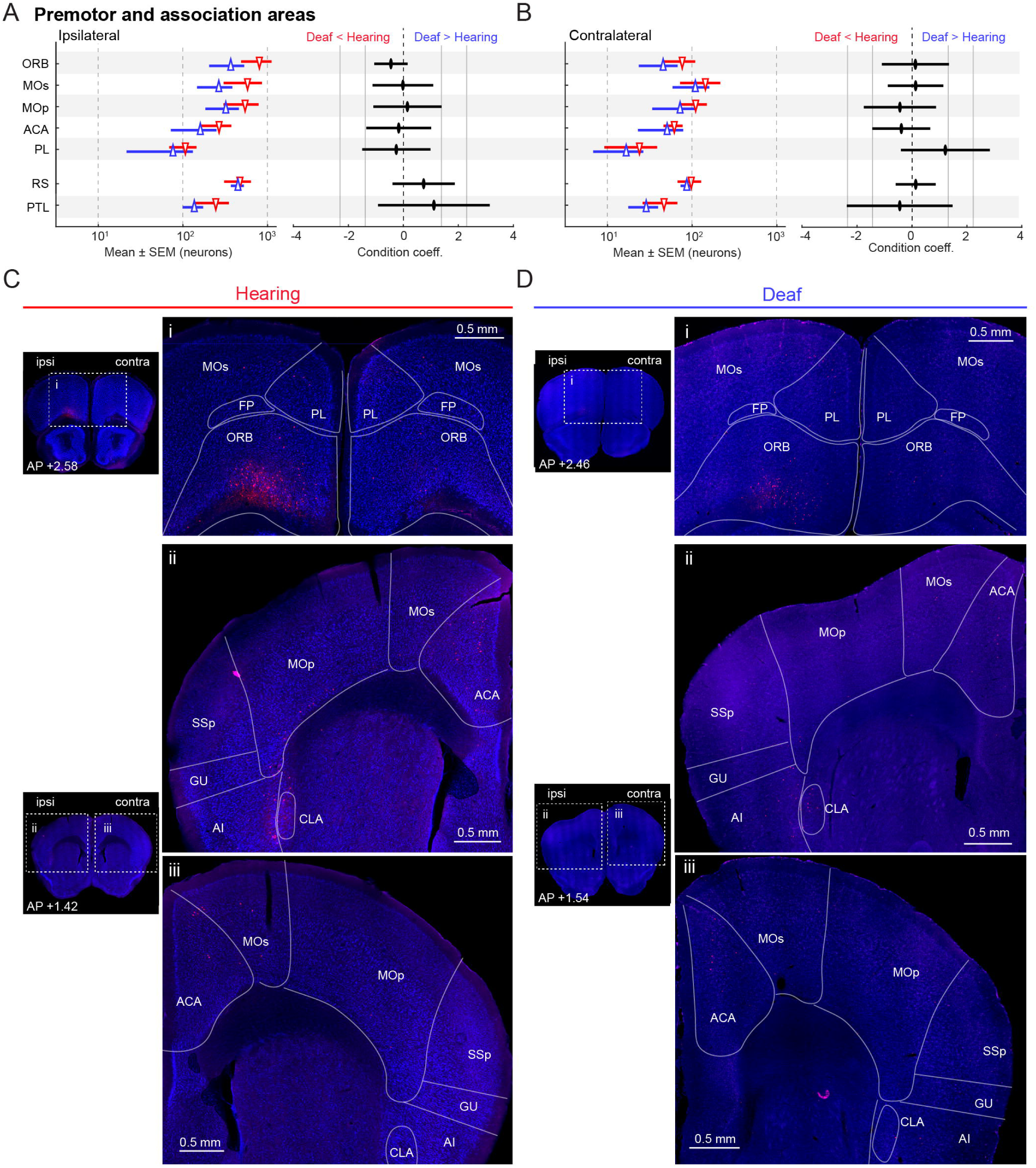
Afferent neuron labeling in premotor and associational cortical areas. **A)** Left, mean ± SEM of auditory cortical afferent neuron counts gathered from ipsilateral premotor (top) or associational (bottom) cortical regions in hearing (red) and deaf (blue) mice. Values plotted on log scale. Right, coefficient and 95% confidence interval of hearing condition for generalized linear model of regional neuron counts. Grey vertical lines, coefficient values corresponding to 4- and 10-fold difference in counts between hearing and deaf mice. **B)** Same as A, but for contralateral regions. **C)** Examples of auditory cortical afferent neuron labeling in premotor cortical regions in a hearing mouse. **D)** Same as C, but for a deaf mouse. Abbreviations: ACA - anterior cingulate cortex; Al - agranular insular cortex; CLA - claustrum; FP - frontal pole; GU - gustatory cortex; MOp - primary motor cortex; MOs - secondary motor cortex; ORB - orbitofrontal cortex; PL - prelimbic cortex; PTL - posterior parietal cortex; RS - retrosplenial cortex.

In several premotor cortical regions, including the orbitofrontal cortex, somatomotor areas, and the anterior cingulate, we observed sparse labeling of afferent neurons in both hearing and deaf mice (Figure 5). Labeling was also evident in associational cortical areas, including the posterior parietal and retrosplenial cortex (example labeling in Figure 4C, 4D, 4G, 4H). In all of these regions, the number of auditory cortical afferent neurons was indistinguishable between hearing and deaf mice. In summary, most auditory cortical afferent neurons are located in the cerebral cortex and, perhaps with the exception of the ipsilateral perirhinal cortex, the number of afferent neurons in all regions of the cortex is unaffected by deafness.

### Auditory experience influences auditory cortical input from the amygdala

We also examined the effects of auditory experience on counts of tdTomato-expressing neurons in subcortical regions. In the cortical subplate, tdTomato-expressing neurons were most numerous in the amygdala. When we examined labeling of neurons in individual amygdalar nuclei, we detected a significant reduction in the number of labeled neurons in the basomedial amygdalar nucleus of deaf mice (p = 0.035; Figure 6A-C). In contrast, counts of labeled neurons from other amygdalar nuclei, including the basolateral amygdala, as well as other regions of the cortical subplate, including the claustrum and endopiriform nucleus, were similar in hearing and deaf mice. Thus, congenital deafness results in selectively diminished input from the basomedial amygdala to the primary auditory cortex.

**Figure 6:**
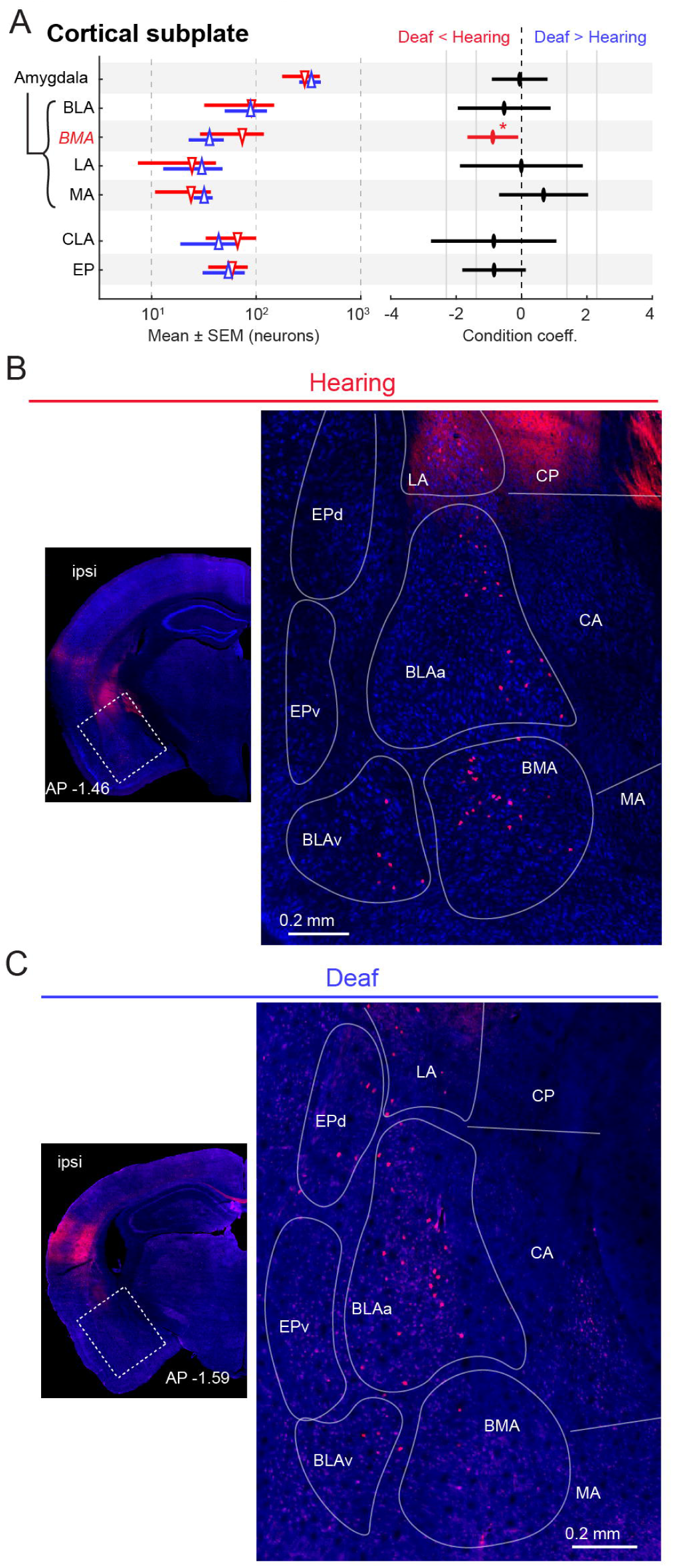
Auditory cortical afferent neuron labeling in the cortical subplate. **A)** Left, mean ± SEM of auditory cortical afferent neuron counts gathered from ipsilateral regions of the cortical subplate in hearing (red) and deaf (blue) mice. Values plotted on log scale. Right, coefficient and 95% confidence interval of hearing condition for generalized linear model of regional neuron counts. Grey vertical lines, coefficient values corresponding to 4- and 10-fold difference in counts between hearing and deaf mice. **B)** Examples of auditory cortical afferent neuron labeling in the cortical subplate of a hearing mouse. **C)** Same as B, but for a deaf mouse. Abbreviations: BLAa and BLAv - anterior and ventral basolateral amygdala; BMA - basomedial amygdala; CA- central amygdala; CLA - claustrum; CP - caudoputamen; EPd and EPv - dorsal and ventral endopiriform nucleus; LA - lateral amygdala; MA - medial amygdala.

### Thalamocortical afferents are markedly reduced in deaf mice

Outside of the cortex, the largest numbers of tdTomato-expressing neurons were located in multiple thalamic nuclei along the anterior-posterior axis. In a subset of nuclei, numbers of labeled neurons were strongly dependent on auditory experience, with a notable reduction in the core of the ventral medial geniculate nucleus and anterior auditory thalamic nuclei of deaf mice. In deaf mice, the number of tdTomato-labeled neurons was similar in the various divisions of the medial geniculate nucleus (MGv, MGd, MGm), the suprageniculate nucleus (SGN), and the posterior limiting nucleus (POL). All thalamic regions with labeling in deaf mice were also labeled in hearing mice, and neither group showed labeling in the visual lateral posterior nucleus, a site of ectopic expansion of auditory cortical afferents in deaf white cats (Barrone et al., 2013; Chabot et al., 2015). Notably, the effects of deafness on the numbers of labeled neurons in the MGv varied along the anterior-posterior axis (Figure 7B-C, 7F-G). Neurons were numerous in the core of the MGv (i.e., anterior to -3.5 mm from bregma) in hearing mice, but reduced in deaf mice. In fact, deafness was strongly predictive of lower neuron counts from the core MGv (p = 0.04). In contrast, at the posterior margin of the MGv, the number of auditory cortical afferent neurons was indistinguishable between hearing and deaf mice. This divergence in thalamocortical input also extended into thalamic regions anterior to the medial geniculate complex. In mice with intact hearing, the anterior region of the thalamus featured labeling in several nuclei surrounding the mammillothalamic tract, with the majority located in the ventral medial nucleus (VM). In deaf mice, we detected a significant reduction in the number of labeled neurons from VM and the surrounding nuclei (p = 0.02; Figure 7A, 7D-E). In summary, congenital deafness resulted in reduced input from a subset of thalamic nuclei that would typically transmit auditory signals to the auditory cortex.

**Figure 7:**
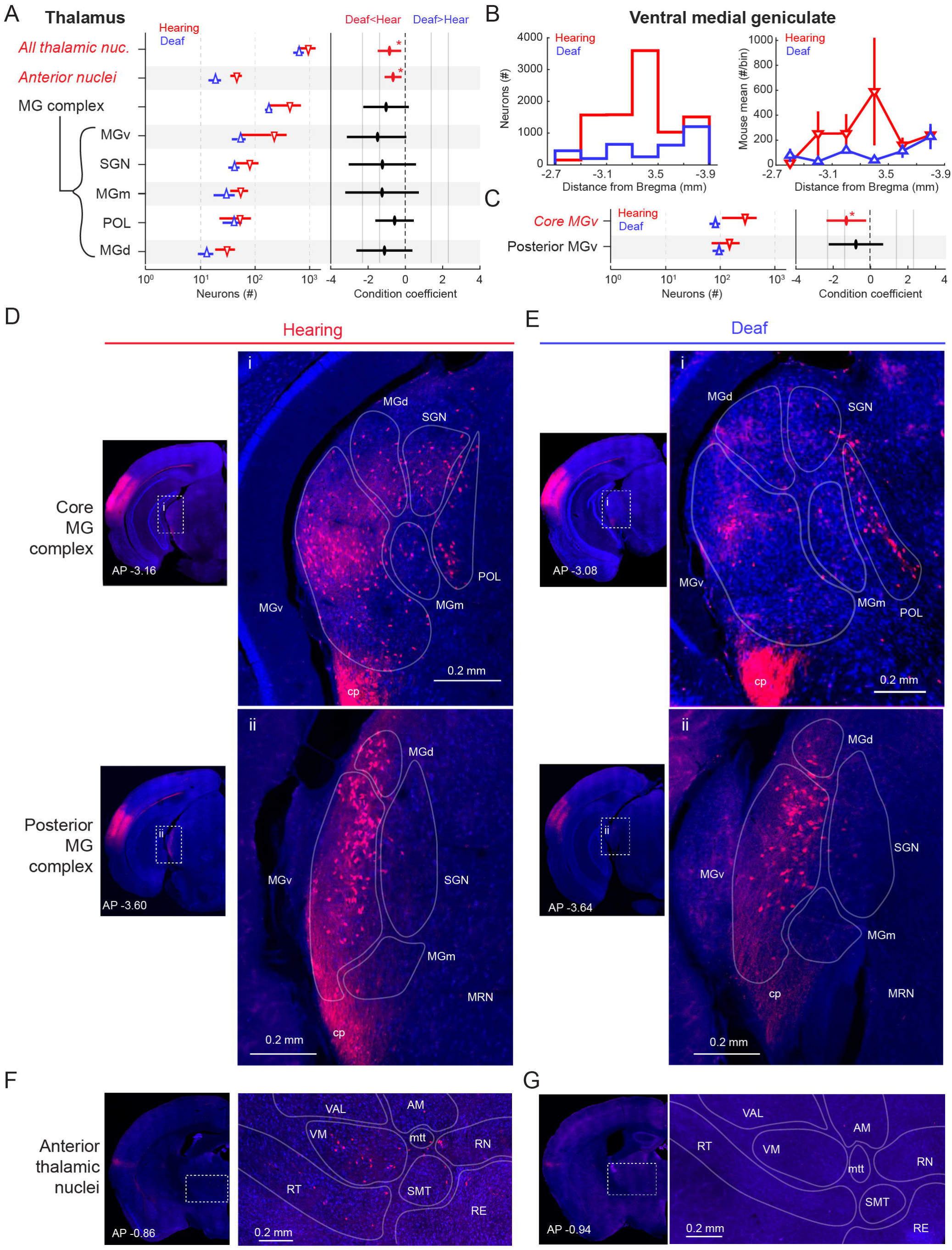
Auditory cortical afferent neuron labeling in the thalamus. **A)** Left, mean ± SEM of afferent neuron counts gathered in entire ipsilateral thalamus, anterior thalamic nuclei, or the medial geniculate and surrounding nuclei (MG complex) in hearing (red) and deaf (blue) mice. Values plotted on log scale. Right, coefficient and 95% confidence interval of hearing condition for generalized linear model of regional neuron counts. Grey vertical lines, coefficient values corresponding to 4- and 10-fold difference in counts between hearing and deaf mice. *, p < 0.05. **B)** Left, histogram of anterior-posterior distribution of all MGv neurons in either hearing (n = 6) or deaf (n=5) mice. Bin = 0.2 mm. Right, mean ± SEM of MGv neuron counts along the anterior-posterior axis in hearing or deaf mice. **C)** same as A, but for neuron counts gathered from the core or posterior regions of the MGv. Core, less than 3.5 mm behind bregma. Posterior, more than 3.5 mm behind bregma. **D)** Example of auditory cortical afferent neuron labeling in the core (i) or posterior (ii) regions of the MG complex in a hearing mouse. **E)** Same as D, but for a deaf mouse. **F)** Example of auditory cortical afferent neuron labeling in the anterior thalamus of a hearing mouse. **G)** Same as F, but for a deaf mouse. Abbreviations - AM - anteromedial nucleus; MGd, MGm, and MGv - dorsal, medial, and ventral part of the medial geniculate nucleus; MRN - midbrain reticular nucleus; mtt - mammillothalamic tract; POL - posterior limiting nucleus; RE - Nucleus of reuniens; RN - rhomboid nucleus; RT - Reticular nucleus; SGN - suprageniculate nucleus; SMT - submedial nucleus; VAL - Ventral anterior-lateral complex; VM - ventromedial nucleus.

## Discussion

Here, we used an intersectional genetic strategy to label, count and compare auditory cortical afferent neurons across the entire brain of hearing and congenitally deaf adult mice with the same genetic background, often from the same litter, and with highly similar non-auditory experiences. Furthermore, our design accounted for additional factors, such as viral efficacy, age, and sex, that could influence neuroanatomical connections to the auditory cortex. This approach revealed that congenitally deaf mice displayed marked reductions in auditory thalamic input to the primary auditory cortex, particularly in the core of the ventral medial geniculate (MGv) and anterior auditory thalamus. We also found that auditory cortical afferent neurons were reduced specifically in the basomedial amygdala in deaf mice. In contrast, cortical inputs to the auditory cortex were largely unaffected by auditory experience. These findings reveal that an absence of hearing both preserves and alters auditory cortical connectivity, which can inform models of experience-dependent reorganization of auditory cortical function and guide interventional therapies that aim to restore auditory function following hearing loss.

### Reduced lemniscal thalamic input in the auditory cortex of deaf mice

In deaf mice, we observed a marked reduction of auditory cortical afferent neurons in the core of the MGv, which provides lemniscal auditory input to the primary auditory cortex. We also observed a greater density of neurons in the granular layer of the auditory cortex in deaf mice, pointing to an adaptation that may help compensate for decreased lemniscal input. Also, the anterior thalamus, including in the ventromedial nucleus (VM), was nearly devoid of auditory cortical afferent neurons. In hearing mice, these neurons provide tonotopic input to layer 1 of the auditory cortex that facilitates frequency discrimination (Zhao et al., 2025). Thus, two separate thalamic inputs that would typically transmit sound frequency information to the auditory cortex are reduced or absent in deaf mice.

By contrast, the lack of hearing did not affect the number of afferent neurons in the posterior MGv, medial and dorsal subdivisions of MG, the posterior limiting nucleus, or the suprageniculate nucleus of the thalamus, all of which are non-lemniscal sources of auditory cortical input. A noteworthy feature is that many neurons in the non-lemniscal pathway respond to both auditory and non-auditory sensory stimuli (Benedek et al., 1997; Bordi & LeDoux, 1994; Lohse et al., 2021). One possibility is that, in the absence of hearing, non-auditory sensory stimulation of these neurons is sufficient to sustain a normal amount of non-lemniscal input to the auditory cortex.

The preservation of auditory cortical afferent neurons in posterior MGv, which projects mainly to the secondary auditory cortex in the mouse (Garcia et al., 2025; Ohga et al., 2018), indicates that retro-Cre viral particles likely spread beyond the boundaries of the primary auditory cortex and into secondary auditory cortical areas. In hearing mice, primary and secondary areas are delineated functionally by imaging sound responses in mesoscale (Garcia et al., 2025; Ohga et al., 2018; Vasquez-Lopez et al., 2017; Wang et al., 2024). The anatomically-defined regional boundaries used here are not necessarily aligned with these functionally-defined boundaries, and delineating auditory cortical subregions with sound responses is not possible in deaf animals. Regardless, our data indicate that lemniscal connections between the core MGv and the auditory cortex require hearing to be established or maintained, whereas non-lemniscal connections are preserved, even in conditions of congenital deafness.

### Neuroanatomical connections from the basomedial amygdala are reduced

In the auditory cortex of deaf mice, we also noted reduced input from the basomedial nucleus of the amygdala (BMA), while other amygdalar nuclei were unaffected. Loss of this projection from the BMA to the auditory cortex might be expected to impact fear and anxiety, two states which the BMA broadly regulates (Adhikari et al., 2015). Consistent with this view, we observed that TMC1^Δ/Δ^ mice were especially docile, even compared with their hearing littermates. The BMA also facilitates social interactions by integrating and relaying multisensory signals to the hypothalamus and cortex (Adamec & Stark-Adamec, 1983; Adhikari et al., 2015; McDonald, 1998; Nowlan et al., 2025). Within this framework, afferents from the BMA to the auditory cortex provide multisensory input to the auditory cortex that modulates responses to sound. For example, afferents from the basal amygdala (i.e., the BMA and basolateral amygdala) transmit mouse pup olfactory cues to the auditory cortex, which, in postpartum or surrogate dams, enhance responses to pup calls (Nowlan et al., 2025). In deaf mice, reduced input from the BMA would be expected to disrupt modulation of the auditory cortex by pup olfactory cues, and perhaps modulation by cues from adult mice as well. In fact, we previously found that a smaller proportion of neurons in the auditory cortex of adult deaf TMC1^Δ/Δ^ mice are modulated by adult female odor (53% in hearing vs. 32% in deaf; Harmon et al., 2024, Supplementary table 1). The reduction of input from the BMA highlights that some adaptations of the auditory cortex that support species-specific social behaviors in hearing animals depend on auditory experience to be established or maintained.

### Connections from motor and non-auditory sensory areas are preserved in deaf mice

The present study confirms that the auditory cortex is a site of remarkable convergence of input from many other cortical regions, including those that encode visual, somatosensory, and movement-related information. One function of these cortico-cortical inputs is to enhance sound encoding in the auditory cortex when auditory stimuli are strongly correlated with other non-auditory stimuli or with movement (Atilgan et al., 2018; Audette et al., 2022; Bigelow et al., 2022; Schneider et al., 2018). However, visual and somatosensory stimuli can also activate a subset of auditory cortical neurons even when presented in the absence of sound (Bigelow et al., 2022; Bizley et al., 2007; Iurilli et al., 2012; Xiao et al., 2022). In the auditory cortex of deaf subjects, these non-auditory sensory responses are enhanced at the population (Auer et al., 2007; Finney et al., 2001; Hunt et al., 2006; Karns et al., 2012) and single unit levels (Allman et al., 2009; Meredith & Allman, 2012). This enhancement seems to improve perceptual capabilities of non-auditory sensory systems, perhaps by co-opting auditory cortical circuits that would typically respond to sound (Lomber et al., 2010; Meredith et al., 2011; Neville & Lawson, 1987). In the genetically deaf white cat and in surgically deafened ferrets, enhanced non-auditory sensory responses are accompanied by increased connectivity of visual and somatosensory cortex with secondary auditory cortical regions (Barone et al., 2013; Kok et al., 2014; Wong et al., 2014). By contrast, input from these areas to the primary auditory cortex is similar in deaf subjects (Barone et al., 2013; Chabot et al., 2015; Meredith & Allman, 2012), in agreement with our findings.

The auditory cortex is also strongly modulated by movements, including vocal gestures (Numminen & Curio, 1999), a property that depends in part on frontal cortical inputs (Nelson et al., 2013; Schneider et al., 2014). The present study establishes that inputs from a variety of frontal cortical regions, including the orbitofrontal cortex and primary and secondary motor cortex, are preserved in congenitally deaf mice. Despite this anatomical preservation, we previously found that the proportion of auditory cortical neurons that are modulated by locomotion (35% in hearing vs. 20% in deaf) or vocalization (62% vs. 24%) is reduced in deaf TMC1^Δ/Δ^ mice, and that movement-related modulation in deaf mice is enhanced in magnitude and altered in timing (Harmon et al., 2024). Taken together, these anatomical and physiological observations support the idea that congenital deafness results in functional reorganization of pathways that convey non-auditory sensory and movement-related signals to the auditory cortex. One possibility is that these forms of adaptation are interdependent, with non-auditory sensory processing expanding in the auditory cortex at the expense of modulation by movement. If so, the persistence of enhanced non-auditory sensory modulation in the auditory cortex following restoration hearing, as has been described in recipients of cochlear implants (Doucet et al., 2006; Lee et al., 2001; Strelnikov et al., 2013), may interfere with movement modulation of the auditory cortex and associated behaviors, including vocalization.

### Potential challenges to restoring auditory cortical sound processing and hearing

The present finding that lemniscal thalamic input to the auditory cortex is markedly reduced in TMC1^Δ/Δ^ mice highlights potential challenges to restoring auditory cortical function (and hearing) in congenitally deaf subjects. One possibility is that the lemniscal pathway regenerates following cochlear implants or gene therapies designed to treat congenital deafness. In fact, following cochlear implant surgery in previously deaf cats or humans, electrical stimulation of the cochlea initially elicits low amplitude, uniform auditory cortical responses that gradually become more dynamic with several months of experience (Klinke et al., 1999; Kral et al., 2002; Kral & Sharma, 2012), a process that may reflect functional recovery of the lemniscal thalamocortical pathway. Alternatively, if these lemniscal connections are permanently lost, then therapies that aim to restore full auditory cortical function must act through other pathways, including non-lemniscal thalamocortical and cortico-cortical projections. Notably, in subjects with normal hearing, the non-lemniscal thalamic pathway is poorly suited for representing the spectral and temporal features of sound with high fidelity (Anderson & Linden, 2011; Garcia et al., 2025; Rodrigues-Dagaeff et al., 1989), and cortico-cortical pathways have been implicated in transmitting visual and motoric information to the auditory cortex, features that may interfere with hearing recovery. Therefore, an important goal of future experiments will be to examine whether the lemniscal thalamocortical pathway recovers anatomically and/or functionally in congenitally deaf subjects that receive hearing restoration therapy, or instead is lost permanently. Mapping of auditory cortical afferents in mouse models where gene therapies can be used to correct congenital deficits in cochlear hair cell function provide a powerful platform for achieving this goal.

## Methods

### *EXPERIMENTAL* MODEL AND STUDY PARTICIPANT DETAILS

#### Animals and husbandry

All surgical and experimental procedures were approved by the Duke University Institutional Animal Care and Use Committee. TMC1^Δ/Δ^ mice (Kawashima et al., 2011, courtesy of Jeffery Holt, Harvard University) were crossed with B6.Cg-Gt(ROSA)26Sortm14(CAG-tdTomato)Hze mice (Ai14; Jackson Labs #7914). Double heterozygous offspring were crossed with TMC1^Δ/Δ^ mice to generate mice that were either TMC1^+/Δ^ (hearing) or TMC1^Δ/Δ^ (deaf) and Ai14^+/-^ or Ai14^-/-^. Mice were genotyped for Ai14 (Transnetyx) and the null TMC1 allele. Hearing was confirmed by measuring auditory brainstem responses to 1 ms click stimuli (>500 presentations), recorded with subdermal low-impendence monopolar electrodes (Technomed, TE/AP 2535), amplified with a microelectrode AC amplifier (AM Systems, Model 1800), and digitized with a Power 1401 data acquisition board at 10 kHz (Cambridge Electronic Design).

### METHOD DETAILS

#### Surgery

Adult (77 - 190 days old, mean = 140 days) female (n = 3 hearing, 3 deaf) and male (n = 3 hearing, 2 deaf) Ai14^+^ mice were anesthetized with isoflurane and injected with meloxicam for analgesia. Hair was removed and topical bupivacaine was applied to the scalp. An incision was made ∼ 2 mm left of the midline, exposing the coronal and lambdoid skull sutures and the left temporal ridge. The region overlying the left auditory was measured stereotactically (AP: −2.8, ML: +4.4 from bregma) and a craniotomy was made with a rotary drill (Dremel, bit size 5). Mice received viral injections of 40 nL of AAVrg-pgk-Cre (Addgene 24593-AAVrg) and AAV-2/9-hsyn-EGFP-WPRE (Addgene 50465-AAV9), combined in equal volumes. We also made 100 nL injections of AAVrg-pgk-Cre only. While we found that this viral volume and titer induced dense labeling around the injection site that made it impossible to count neurons accurately, sections from the brains of these mice (n = 5 hearing, 5 deaf) were used for analysis of auditory cortical cytoarchitecture. The tip of a glass pipette loaded with virus was lowered to 600 μm below the surface of the brain and 10 nL of virus was pressure injected (Nanoject III, Drummond Scientific; rate = 1 nL/s). To ensure that virus was injected into all layers of the auditory cortex, the procedure was repeated three times after retracting 100 μm per injection, resulting in 10 nL viral injections at 600, 500, 400 and 300 μm from the brain surface. The craniotomy was covered with bone wax (Ethicon), the incision was sutured closed, and the mouse was returned to its home cage to recover. Meloxicam was injected once more ∼12 hrs after the surgery.

#### Histology

Histology and quantification was performed by researchers who were blinded to the mouse’s hearing status. Mice were euthanized by transcardial perfusion 14-21 days after injection of viral vectors. Brains were fixed in 4% paraformaldehyde (PFA) in phosphate buffer solution for 24 hrs, transferred to a 30% sucrose PFA solution until saturation, embedded in Tissue Tek O.C.T. compound in dry ice, and sliced into 50-µm coronal sections on a Leica CM1950 cryostat.

Sections were incubated with blue fluorescent Nissl stain (Neurotrace 435/455) and mounted on glass slides. Images of tdTomato, enhanced GFP, and blue Nissl fluorescence were acquired as a tile scan from every fourth section with a Zeiss 710 inverted confocal microscope in the Duke University Light Microscopy Core Facility. The green channel was omitted when no somatic signal was present. The magnification, laser power, aperture, pixel binning, digital amplification, and tile scan overlap and stitching were matched across sections and mice.

### QUANTIFICATION AND STATISTICAL ANALYSIS

#### Cell density and cytoarchitecture analysis

Measurements of the relative cell density and cortical depth were made using custom MATLAB (2023) code. For each mouse, sections that included the full depth of the primary auditory cortex in the hemisphere contralateral to the injection site were identified. Sections in which the auditory cortex was damaged, occluded, or stained unevenly were excluded. For measurements of estimated regional cell density, we divided the mean amplitude of pixels within the auditory cortex by the mean amplitude of the cortical area dorsal to the auditory cortex. For measurements of cortical depth, we used the *drawline* command to insert a vector from the inferior to superior surface of the auditory cortex in each section and measured its length. To measure cell density through the depth of the cortex, we interpolated 41 points along the vector, measured the mean amplitude of pixels that fell within the area defined by consecutive points, then calculated the z score using the mean and standard deviation of the signal amplitude in the cortical region dorsal to the auditory cortex. For measurements of regional cell density and cortical depth, statistically significant differences between hearing and deaf mice were tested with unpaired students t-tests. Differences in cell density across the depth of the cortex were tested with a repeated measures ANOVA and post hoc Tukey honest statistical difference test.

#### Image processing and quantification

Confocal images were downsampled to half their original dimensions, matched for orientation, and split by color channel into three separate images. Fluorescent neurons were identified, counted, and assigned to a brain region with the QUINT workflow (Yates et al., 2019). Using QuickNii software, Nissl stain images were aligned to projections from the Allen Institute Mouse Brain Atlas (2017), which were manually adjusted to match the precise sectioning angle. We noted that atlas projections were often poorly aligned for posterior sections in which the cortical hemispheres and the brainstem are not connected. Therefore, after generating atlas projections for the ipsilateral cortical hemisphere, we also generated projections that were aligned to the ipsilateral thalamus and contralateral cortex but that maintained the same anterior-posterior position and rotation.

Fluorescent neurons were identified in tdTomato and GFP images using the Pixel and Object Classification workflow in ilastik. The pixel classifier was manually trained with several example images to distinguish fluorescent and non-fluorescent pixels. The output of the pixel classifier fed into the object classifier, which identified ROIs (i.e., neuron cell bodies) through hysteresis. Object segmentation was optimized to identify as many somatic ROIs as possible while limiting merged objects by systematically adjusting smoothing and threshold values.

Binary masks of cell bodies were imported into NUTIL software, which calculated the center position and area of each ROI in pixel space, combined anatomical atlas projections and mask images, and assigned each ROI to a brain region. Data from NUTIL and anterior-posterior positions of sections relative to bregma were imported into MATLAB. Each ROI was designated as ipsi- or contralateral to the injection site, and three dimensional coordinates of each ROI’s position relative to bregma were calculated.

The injection site was defined as the region with the most GFP labeled ROIs. Brains with injections centered outside of the primary auditory cortex were excluded. Coordinates of the injection site were calculated as the mean position of all GFP ROIs, weighted by ROI size. Some sections, particularly those proximal to the injection site, featured ROIs comprising multiple merged neurons. To estimate the number of labeled neurons within an ROI, the ROI pixel area was divided by the modal pixel area calculated across all ROIs (26 pixels) and rounded to the nearest integer. We noted several instances in which fluorescence from axon terminal fields of corticothalamic projection neurons was classified as somatic ROIs in the thalamus. Therefore, we manually identified thalamic ROIs and generated binary mask images which were used to count auditory cortical afferent neurons in the thalamus after assigning them to thalamic nuclei.

#### Generalized multivariate linear regression models of regional neuron counts

To estimate the effect of deafness on regional afferent neuron counts, the number of tdTomato-labeled neurons within each region was modeled in MATLAB as Poisson distribution using the *fitglm* command. The number of GFP-labeled neurons throughout the entire brain, the number of tdTomato-labeled neurons outside of the target region, viral incubation time, the mouse’s age at the time of injection, sex, and hearing status were tabulated and designated as fixed effect predictor variables, while mouse identity was designated as random effect variable. The natural log of region area was included as an offset normalization term. To be considered for the analysis, the mean neuron count from either hearing or deaf mice had to surpass five neurons, a criterion that was satisfied by 26 cortical regions (both hemispheres), seven thalamic regions, and six cortical subplate regions. The values and 95% confidence intervals of the weighting coefficient of hearing condition are reported in figures 3-8. For models that returned a significant effect of hearing condition, the raw p-value of the hearing condition coefficient is reported in the text. Bayesian inference criterion values were, on average, lower for the full models than for models constructed only from the number of GFP neurons, remaining tdTomato neurons, and hearing condition (35.1 vs. 36.5), indicating that inclusion of additional predictor variables improved model performance.

**Table 1:** Brain region abbreviations.

| **Abbreviation** | **Full Brain Region Name** |
| --- | --- |
| ACA | Anterior cingulate area |
| AI | Agranular insular area |
| AM | Anteromedial nucleus of the thalamus |
| AUDd | Dorsal auditory area |
| AUDp | Primary auditory area |
| AUDpos | Posterior auditory area |
| AUDv | Ventral auditory area |
| BLA | Basolateral amygdalar nucleus |
| BLAa | Basolateral amygdalar nucleus, anterior part |
| BLAv | Basolateral amygdalar nucleus, ventral part |
| BMA | Basomedial amygdalar nucleus |
| CEA | Central amygdalar nucleus |
| cc | Corpus callosum |
| CLA | Clastrum |
| CP | Caudoputamen |
| cp | Cerebral peduncle |
| ECT | Ectorhinal area |
| ECTlat | Ectorhinal area, lateral part |
| ENT | Entorhinal area |
| EPd | Endopiriform nucleus, dorsal part |
| EPv | Endopiriform nucleus, ventral part |
| FP | Frontal pole, cerebral cortex |
| GU | Gustatory areas |
| HPF | Hippocampal formation |
| LA | Lateral amygdalar nucleus |
| MA | Medial amygdalar nucleus |
| MGd | Medial geniculate complex, dorsal part |
| MGm | Medial geniculate complex, medial part |
| MGv | Medial geniculate complex, ventral part |
| MOp | Primary motor area |
| MOs | Secondary motor area |
| MRN | Midbrain reticular nucleus |
| mtt | Mammillothalamic tract |
| ORB | Orbital area |
| PIR | Piriform area |
| PL | Prelimbic area |
| POL | Posterior limiting nucleus of the thalamus |
| RE | Nucleus of reuniens |
| RH | Rhomboid nucleus of the thalamus |
| RS | Retrosplenial area |
| RT | Reticular nucleus of the thalamus |
| SGN | Suprageniculate nucleus |
| SMT | Submedial nucleus of the thalamus |
| SSp | Primary somatosensory area |
| SSs | Supplemental somatosensory area |
| TEa | Temporal association area |
| VAL | Ventral anterior-lateral complex of the thalamus |
| VISal | Anterolateral visual area |
| VISam | Anteromedial visual area |
| VISlat | Lateral visual area |
| VISp | Primary visual area |
| VISpl | Posterolateral visual area |
| VISpm | Posteromedial visual area |
| VM | Ventromedial nucleus of the thalamus |

## References

Adamec, R. E., & Stark-Adamec, C. I. (1983). Limbic control of aggression in the cat. Progress in Neuro-Psychopharmacology & Biological Psychiatry, 7(4-6), 505–512.

Adhikari, A., Lerner, T. N., Finkelstein, J., Pak, S., Jennings, J. H., Davidson, T. J., Ferenczi, E., Gunaydin, L. A., Mirzabekov, J. J., Ye, L., Kim, S.-Y., Lei, A., & Deisseroth, K. (2015). Basomedial amygdala mediates top-down control of anxiety and fear. Nature, 527(7577), 179–185.

Akil, O., Seal, R. P., Burke, K., Wang, C., Alemi, A., During, M., Edwards, R. H., & Lustig, L. R. (2012). Restoration of hearing in the VGLUT3 knockout mouse using virally mediated gene therapy. Neuron, 75(2), 283–293.

Allman, B. L., Keniston, L. P., & Meredith, M. A. (2009). Adult deafness induces somatosensory conversion of ferret auditory cortex. Proceedings of the National Academy of Sciences of the United States of America, 106(14), 5925–5930.

Anderson, L. A., & Linden, J. F. (2011). Physiological differences between histologically defined subdivisions in the mouse auditory thalamus. Hearing Research, 274(1-2), 48–60.

Atilgan, H., Town, S. M., Wood, K. C., Jones, G. P., Maddox, R. K., Lee, A. K. C., & Bizley, J. K. (2018). Integration of Visual Information in Auditory Cortex Promotes Auditory Scene Analysis through Multisensory Binding. Neuron, 97(3), 640–655.e4.

Audette, N. J., Zhou, W., La Chioma, A., & Schneider, D. M. (2022). Precise movement-based predictions in the mouse auditory cortex. Current Biology : CB, 32(22), 4925–4940.e6.

Auer, E. T., Jr, Bernstein, L. E., Sungkarat, W., & Singh, M. (2007). Vibrotactile activation of the auditory cortices in deaf versus hearing adults. Neuroreport, 18(7), 645–648.

Barone, P., Lacassagne, L., & Kral, A. (2013). Reorganization of the connectivity of cortical field DZ in congenitally deaf cat. PloS One, 8(4), e60093.

Benedek, G., Perény, J., Kovács, G., Fischer-Szátmári, L., & Katoh, Y. Y. (1997). Visual, somatosensory, auditory and nociceptive modality properties in the feline suprageniculate nucleus. Neuroscience, 78(1), 179–189.

Berger, C., Kühne, D., Scheper, V., & Kral, A. (2017). Congenital deafness affects deep layers in primary and secondary auditory cortex. The Journal of Comparative Neurology, 525(14), 3110–3125.

Bigelow, J., Morrill, R. J., Olsen, T., & Hasenstaub, A. R. (2022). Visual modulation of firing and spectrotemporal receptive fields in mouse auditory cortex. Current Research in Neurobiology, 3, 100040.

Bizley, J. K., Nodal, F. R., Bajo, V. M., Nelken, I., & King, A. J. (2007). Physiological and anatomical evidence for multisensory interactions in auditory cortex. *Cerebral Cortex (New York*, N.Y*. :* 1991), 17(9), 2172–2189.

Bordi, F., & LeDoux, J. E. (1994). Response properties of single units in areas of rat auditory thalamus that project to the amygdala. II. Cells receiving convergent auditory and somatosensory inputs and cells antidromically activated by amygdala stimulation. Experimental Brain Research, 98(2), 275–286.

Chabot, N., Butler, B. E., & Lomber, S. G. (2015). Differential Modification of Cortical and Thalamic Projections to Cat Primary Auditory Cortex Following Early- and Late-Onset Deafness. The Journal of Comparative Neurology, 523(15), 2297–2320.

da Costa, N. M., Martin, K. A. C., & Sägesser, F. D. (2017). A weighted graph of the projections to mouse auditory cortex. In bioRxiv. bioRxiv. 10.1101/228726

Doucet, M. E., Bergeron, F., Lassonde, M., Ferron, P., & Lepore, F. (2006). Cross-modal reorganization and speech perception in cochlear implant users. Brain : A Journal of Neurology, 129(Pt 12), 3376–3383.

Eliades, S. J., & Wang, X. (2003). Sensory-motor interaction in the primate auditory cortex during self-initiated vocalizations. Journal of Neurophysiology, 89(4), 2194–2207.

Finney, E. M., Fine, I., & Dobkins, K. R. (2001). Visual stimuli activate auditory cortex in the deaf. Nature Neuroscience, 4(12), 1171–1173.

Francis, N. A., Winkowski, D. E., Sheikhattar, A., Armengol, K., Babadi, B., & Kanold, P. O. (2018). Small Networks Encode Decision-Making in Primary Auditory Cortex. Neuron, 97(4), 885–897.e6.

Gale, D. J., Areshenkoff, C. N., Honda, C., Johnsrude, I. S., Flanagan, J. R., & Gallivan, J. P. (2021). Motor Planning Modulates Neural Activity Patterns in Early Human Auditory Cortex. *Cerebral Cortex (New York*, N.Y*. :* 1991), 31(6), 2952–2967.

Garcia, M. M., Kline, A. M., Onodera, K., Tsukano, H., Dandu, P. R., Acosta, H. C., Kasten, M. R., Manis, P. B., & Kato, H. K. (2025). Noncanonical short-latency auditory pathway directly activates deep cortical layers. Nature Communications, 16(1), 5911.

Harmon, T. C., Madlon-Kay, S., Pearson, J., & Mooney, R. (2024). Vocalization modulates the mouse auditory cortex even in the absence of hearing. Cell Reports, 43(8), 114611.

Hunt, D. L., Yamoah, E. N., & Krubitzer, L. (2006). Multisensory plasticity in congenitally deaf mice: how are cortical areas functionally specified? Neuroscience, 139(4), 1507–1524.

Iurilli, G., Ghezzi, D., Olcese, U., Lassi, G., Nazzaro, C., Tonini, R., Tucci, V., Benfenati, F., & Medini, P. (2012). Sound-driven synaptic inhibition in primary visual cortex. Neuron, 73(4), 814–828.

Karns, C. M., Dow, M. W., & Neville, H. J. (2012). Altered cross-modal processing in the primary auditory cortex of congenitally deaf adults: a visual-somatosensory fMRI study with a double-flash illusion. The Journal of Neuroscience : The Official Journal of the Society for Neuroscience, 32(28), 9626–9638.

Kawashima, Y., Géléoc, G. S. G., Kurima, K., Labay, V., Lelli, A., Asai, Y., Makishima, T., Wu, D. K., Della Santina, C. C., Holt, J. R., & Griffith, A. J. (2011). Mechanotransduction in mouse inner ear hair cells requires transmembrane channel-like genes. The Journal of Clinical Investigation, 121(12), 4796–4809.

Klinke, R., Kral, A., Heid, S., Tillein, J., & Hartmann, R. (1999). Recruitment of the auditory cortex in congenitally deaf cats by long-term cochlear electrostimulation. *Science (New York*, N.Y.), 285(5434), 1729–1733.

Kok, M. A., Chabot, N., & Lomber, S. G. (2014). Cross-modal reorganization of cortical afferents to dorsal auditory cortex following early- and late-onset deafness. The Journal of Comparative Neurology, 522(3), 654–675.

Kral, A., Hartmann, R., Tillein, J., Heid, S., & Klinke, R. (2002). Hearing after congenital deafness: central auditory plasticity and sensory deprivation. *Cerebral Cortex (New York*, N.Y*. :* 1991), 12(8), 797–807.

Kral, A., Schröder, J.-H., Klinke, R., & Engel, A. K. (2003). Absence of cross-modal reorganization in the primary auditory cortex of congenitally deaf cats. Experimental Brain Research, 153(4), 605–613.

Kral, A., & Sharma, A. (2012). Developmental neuroplasticity after cochlear implantation. Trends in Neurosciences, 35(2), 111–122.

Lee, D. S., Lee, J. S., Oh, S. H., Kim, S. K., Kim, J. W., Chung, J. K., Lee, M. C., & Kim, C. S. (2001). Cross-modal plasticity and cochlear implants. Nature, 409(6817), 149–150.

Lohse, M., Dahmen, J. C., Bajo, V. M., & King, A. J. (2021). Subcortical circuits mediate communication between primary sensory cortical areas in mice. Nature Communications, 12(1), 3916.

Lomber, S. G., Meredith, M. A., & Kral, A. (2010). Cross-modal plasticity in specific auditory cortices underlies visual compensations in the deaf. Nature Neuroscience, 13(11), 1421–1427.

McDonald, A. J. (1998). Cortical pathways to the mammalian amygdala. Progress in Neurobiology, 55(3), 257–332.

Meredith, M. A., & Allman, B. L. (2012). Early hearing-impairment results in crossmodal reorganization of ferret core auditory cortex. Neural Plasticity, 2012, 601591.

Meredith, M. A., Kryklywy, J., McMillan, A. J., Malhotra, S., Lum-Tai, R., & Lomber, S. G. (2011). Crossmodal reorganization in the early deaf switches sensory, but not behavioral roles of auditory cortex. Proceedings of the National Academy of Sciences of the United States of America, 108(21), 8856–8861.

Müller-Preuss, P., & Ploog, D. (1981). Inhibition of auditory cortical neurons during phonation. Brain Research, 215(1-2), 61–76.

Napoli, J. L., Camalier, C. R., Brown, A.-L., Jacobs, J., Mishkin, M. M., & Averbeck, B. B. (2021). Correlates of Auditory Decision-Making in Prefrontal, Auditory, and Basal Lateral Amygdala Cortical Areas. The Journal of Neuroscience : The Official Journal of the Society for Neuroscience, 41(6), 1301–1316.

Nelson, A., Schneider, D. M., Takatoh, J., Sakurai, K., Wang, F., & Mooney, R. (2013). A circuit for motor cortical modulation of auditory cortical activity. The Journal of Neuroscience : The Official Journal of the Society for Neuroscience, 33(36), 14342–14353.

Neville, H. J., & Lawson, D. (1987). Attention to central and peripheral visual space in a movement detection task. III. Separate effects of auditory deprivation and acquisition of a visual language. Brain Research, 405(2), 284–294.

Nist-Lund, C. A., Pan, B., Patterson, A., Asai, Y., Chen, T., Zhou, W., Zhu, H., Romero, S., Resnik, J., Polley, D. B., Géléoc, G. S., & Holt, J. R. (2019). Improved TMC1 gene therapy restores hearing and balance in mice with genetic inner ear disorders. Nature Communications, 10(1), 236.

Nowlan, A. C., Choe, J., Tromblee, H., Kelahan, C., Hellevik, K., & Shea, S. D. (2025). Multisensory integration of social signals by a pathway from the basal amygdala to the auditory cortex in maternal mice. Current Biology : CB, 35(1), 36–49.e4.

Numminen, J., & Curio, G. (1999). Differential effects of overt, covert and replayed speech on vowel-evoked responses of the human auditory cortex. Neuroscience Letters, 272(1), 29–32.

Ohga, S., Tsukano, H., Horie, M., Terashima, H., Nishio, N., Kubota, Y., Takahashi, K., Hishida, R., Takebayashi, H., & Shibuki, K. (2018). Direct Relay Pathways from Lemniscal Auditory Thalamus to Secondary Auditory Field in Mice. *Cerebral Cortex (New York*, N.Y*. :* 1991), *28*(12), 4424–4439.

Pan, J., Yang, C., Li, R., Li, J., Sun, P., Liu, Y., Liu, K., Liao, X., Jia, H., Yu, Z., Chen, X., & Wang, M. (2025). Brain-wide atlas of outputs and inputs of mouse auditory cortex revealed by modified viral labeling strategies. Fundamental Research. 10.1016/j.fmre.2025.08.014

Rodrigues-Dagaeff, C., Simm, G., De Ribaupierre, Y., Villa, A., De Ribaupierre, F., & Rouiller, E. M. (1989). Functional organization of the ventral division of the medial geniculate body of the cat: evidence for a rostro-caudal gradient of response properties and cortical projections. Hearing Research, 39(1-2), 103–125.

Roux, I., Safieddine, S., Nouvian, R., Grati, M. ’hamed, Simmler, M.-C., Bahloul, A., Perfettini, I., Le Gall, M., Rostaing, P., Hamard, G., Triller, A., Avan, P., Moser, T., & Petit, C. (2006). Otoferlin, defective in a human deafness form, is essential for exocytosis at the auditory ribbon synapse. Cell, 127(2), 277–289.

Schneider, D. M., Nelson, A., & Mooney, R. (2014). A synaptic and circuit basis for corollary discharge in the auditory cortex. Nature, 513(7517), 189–194.

Schneider, D. M., Sundararajan, J., & Mooney, R. (2018). A cortical filter that learns to suppress the acoustic consequences of movement. Nature, 561(7723), 391–395.

Strelnikov, K., Rouger, J., Demonet, J.-F., Lagleyre, S., Fraysse, B., Deguine, O., & Barone, P. (2013). Visual activity predicts auditory recovery from deafness after adult cochlear implantation. Brain : A Journal of Neurology, 136(Pt 12), 3682–3695.

Vasquez-Lopez, S. A., Weissenberger, Y., Lohse, M., Keating, P., King, A. J., & Dahmen, J. C. (2017). Thalamic input to auditory cortex is locally heterogeneous but globally tonotopic. eLife, 6. 10.7554/eLife.25141

Wang, C., Jiang, Z.-Y., Chai, J.-Y., Chen, H.-S., Liu, L.-X., Dang, T., & Meng, X.-M. (2024). Mouse auditory cortex sub-fields receive neuronal projections from MGB subdivisions independently. Scientific Reports, 14(1), 7078.

Wong, C., Chabot, N., Kok, M. A., & Lomber, S. G. (2014). Modified areal cartography in auditory cortex following early- and late-onset deafness. *Cerebral Cortex (New York*, N.Y*. :* 1991), *24*(7), 1778–1792.

Xiao, Y.-J., Wang, L., Liu, Y.-Z., Chen, J., Zhang, H., Gao, Y., He, H., Zhao, Z., & Wang, Z. (2022). Excitatory Crossmodal Input to a Widespread Population of Primary Sensory Cortical Neurons. Neuroscience Bulletin, 38(10), 1139–1152.

Yates, S. C., Groeneboom, N. E., Coello, C., Lichtenthaler, S. F., Kuhn, P.-H., Demuth, H.-U., Hartlage-Rübsamen, M., Roßner, S., Leergaard, T., Kreshuk, A., Puchades, M. A., & Bjaalie, J. G. (2019). QUINT: Workflow for Quantification and Spatial Analysis of Features in Histological Images From Rodent Brain. Frontiers in Neuroinformatics, 13, 75.

Zhao, Z., Tang, X., Chen, Y., Tao, J., Polat, M., Yang, Z., Yang, L., Wang, M., Liang, S., Zhang, K., Zhang, Y., Zhang, C., Wang, L., Wang, Y., Konnerth, A., Jia, H., Xiong, W., Liao, X., Li, S. C., & Chen, X. (2025). A parallel tonotopically arranged thalamocortical circuit for sound processing. Neuron, 113(12), 1998–2013.e6.

